# An estimator of the Opportunity for Selection that is independent of mean fitness

**DOI:** 10.1101/2020.05.06.081224

**Authors:** Robin S. Waples

## Abstract

Variation among individuals in number of offspring (fitness, *k*) sets an upper limit to the evolutionary response to selection. This constraint is quantified by Crow’s Opportunity for Selection (*I*), which is the variance in relative fitness 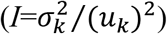. Crow’s *I* has been widely used but remains controversial because it depends on mean offspring number in a sample 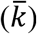. Here I used a generalized Wright-Fisher model that allows for unequal probabilities of producing offspring to evaluate behavior of Crow’s *I* and related indices under a wide range of sampling scenarios. Analytical and numerical results are congruent and show that rescaling the sample variance 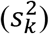 to its expected value at a fixed 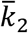 removes dependence of *I* on mean offspring number, but the result still depends on choice of 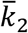. A new index is introduced, 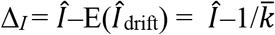, which makes *Î* independent of sample 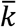 without the need for variance rescaling. Δ_*I*_ has a straightforward interpretation as the component of variance in relative fitness that exceeds that expected under a null model of random reproductive success. Δ_*I*_ can be used to directly compare estimates of the Opportunity for Selection for samples from different studies, different sexes, and different life stages.

---

Over a half century ago, James Crow introduced two indices that quantify variation in reproductive success among individuals, based on the mean (*μ_k_*) and variance 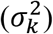 in the number of offspring per individual (*k*). Crow and Morton (1955) defined the ratio 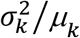 as the Index of Variability, denoted here by φ. With random reproductive success of *N* parents, the expected variance in offspring number due to drift 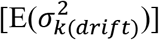 takes the form of a binomial variance *μ_k_*(*N*-1)/*N*, which is often approximated by the Poisson variance 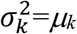, leading to E(φ_drift_) ≈ 1. Therefore, the Index of Variability is a useful indicator of the degree to which variance in offspring number is overdispersed, relative to the Poisson expectation, which in turn has a strong influence on the effective population size (*N_e_*) and the key ratio *N_e_*/*N*. Shortly thereafter, Crow (1958) defined the parameter 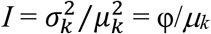, which he called the Index of Total Selection, but subsequent authors generally refer to this as the Opportunity for Selection (Arnold and Wade 1984a, b; Clutton-Brock 1988; Brodie et al. 1995). Crow’s *I* measures “the total amount of selection possible, given the demographics of a population. It answers the question, By what fraction would the mean population fitness increase in one generation of selection if its heritability were perfect (i.e., *h*^2^ = 1)?” (Crow 1989, p 776). The Opportunity for Selection thus sets an upper limit to the evolutionary response to selection; mathematically, *I* also represents the variance in relative fitness (Walsh and Lynch 2018).

Crow and Morton (1955) showed that the Index of Variability is very sensitive to mean offspring number (= mean fitness) in a sample 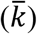, and it is well known that Crow’s *I* also depends on 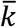. In a stable population of diploids, each parent contributes on average half the genes to two adult offspring; therefore, assuming exhaustive sampling of offspring, *μ_k_* = 2. However, *μ_k_*>2 is possible for increasing populations, and 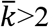 can easily occur when sampling juvenile offspring of a highly fecund species (e.g., a marine fish). Conversely, 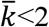 is possible for declining populations or when only a portion of the offspring are sampled. Therefore, sampling from real populations in nature can produce a wide range of sample 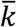 values, even when the population is stable. Dependence of the estimators 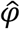 and *Î* on 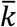 means that, based on experimental design and sampling effort, the same underlying biological process could lead to very different conclusions about the distribution of reproductive success and the Opportunity for Selection. For this reason, a number of authors have questioned the practical utility of Crow’s *I* for drawing inferences about natural selection (Trail 1985; Downhower et al. 1987; Fairbairn and Wilby 2001).

This topic merits a new look, for two reasons. First, Crow and Morton (1955) proposed a simple solution to the problem caused by dependence of the Index of Variability on 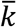: one can rescale 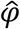 to its expected value at a different mean offspring number using the following equation:

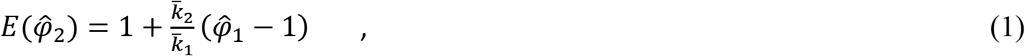

where 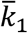 is mean offspring number in the initial sample, 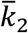 is the target mean offspring number, 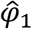 is the initial estimate of the Index of Variability, and 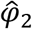 is the rescaled index. Equation 1 solves for the expected value of 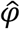, assuming random mortality of offspring until the target 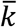 is reached. Allowing for random mortality of individuals in a sample is statistically equivalent to randomly sampling fewer offspring in the first place. Setting 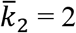 yields the expected value of 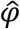 for a constant population. Waples (2002) showed that Equation 1 also can be used for the common situation where 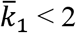; in this case, the result is the value of 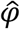 one would expect to find if the same offspring distribution were sampled more intensively, until 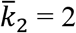. There is nothing magical about scaling to 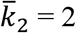, but it is a useful reference point because most populations that persist for any period of time must have a long-term mean offspring number close to 2.

Because of the simple relationship between *I* and φ, Crow and Morton’s method for rescaling φ can also be applied to *I* to remove the dependence on the sample mean. However, few published papers cite both Crow (1958) and Crow and Morton (1955). Crow himself did not mention the 1955 variance-rescaling method in either his original 1958 paper introducing the Index of Total Selection or his 1989 retrospective. Others (e.g., Wade and Arnold 1980 and Cabana and Kramer 1991) noted Crow and Morton’s method but did not use it to rescale *I* values. Therefore, it is important to evaluate the degree to which rescaling raw *I* values can resolve ongoing controversies regarding the Opportunity for Selection.

The second interesting issue is that, under random reproductive success, the Opportunity for Selection has a simple expectation: 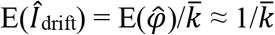. Downhower et al. (1987) suggested, but did not recommend, that the difference between empirical *Î* and 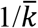 might be used to compare *Î* values from different studies. Their logic was that the result would still be a ratio, and ratios have some undesirable statistical properties. As demonstrated below, Downhower et al.’s idea has some real merit: Although both empirical *Î* and E(*Î*_drift_) depend on sample 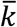, their difference does not and hence is independent of mean fitness in the sample.

Here, analytical and numerical methods are used to explore the relationship between Crow’s Index of Variability and the Opportunity for Selection. Objectives are to: (1) Illustrate the effects of rescaling estimators 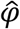 and *Î* to a constant 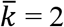, under three different reproductive systems: random reproductive success, and moderate and strong reproductive skew. (2) Introduce a new index Δ_I_ = *Î* - E(*Î*_drift_) and show that this removes the dependence of *Î* on mean offspring number, without any need for rescaling to constant population size. Δ_I_ represents the component of variance in relative fitness that exceeds that due to random demographic stochasticity. (3) Examine an index of relative resource monopolization (*Q*) proposed by Ruzzante et al. (1996) for evaluating the Opportunity for Selection, but recast the index in terms of *I* rather than the variance in offspring number. (4) Use raw and scaled reproductive success data to calculate inbreeding and variance effective sizes and evaluate their sensitivity to the scaling issues that affect φ.

## Methods

Both of the “1”s in Crow and Morton’s Equation 1 are a consequence of using the Poisson approximation that E(φ_drift_) ≈ 1, rather than the exact E(φ_drift_) = (*N*-1)/*N*. The difference is generally negligible, but it can become important when both *N* and mean offspring number are small. The analytical treatment below follows Crow and Morton in using the Poisson approximation. In rescaling empirical data, the following adjustment to Equation 1 can be used to incorporate the exact variance:

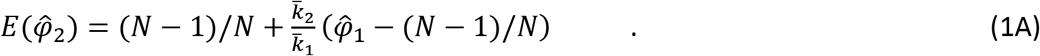

### Analytical model

The model used here is a generalization of the standard Wright-Fisher model of reproduction, whereby *N* monoecious adults each contribute equally to a large (~ infinite) pool of gametes, which unite at random to form the next generation. The current model differs in two respects. First, separate sexes are allowed. For simplicity, the treatment below assumes the sex ratio is equal; if not, the only major difference is that mean offspring number to produce a stable population will be <2 for the more numerous sex and >2 for the less numerous sex. Second, and more importantly, the model allows for unequal contributions to the initial gamete pool, specified by the vector of individual weights *W* = *w*_1_, *w*_2_,… *w_n_* (Figure 1). A version of this weighted Wright-Fisher model was first introduced by Robertson (1961) for the special case of full-sibling families, but the more general treatment here follows Felsenstein (2019). The number of gametes contributed by the *i*^th^ parent (*k_i_*) is proportional to its weight: *k_iα_* = *Dw_i_*, where *i* = 1 … *N, D* is a (very) large constant, and the subscript α indicates that the *k_i_* values apply to the initial gamete pool. It follows that the mean number of gametes per parent in the gamete pool is given by

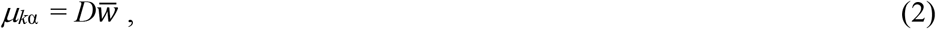

and the variance of initial *k_i_* is

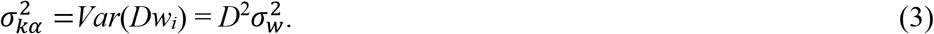

**Figure 1.**
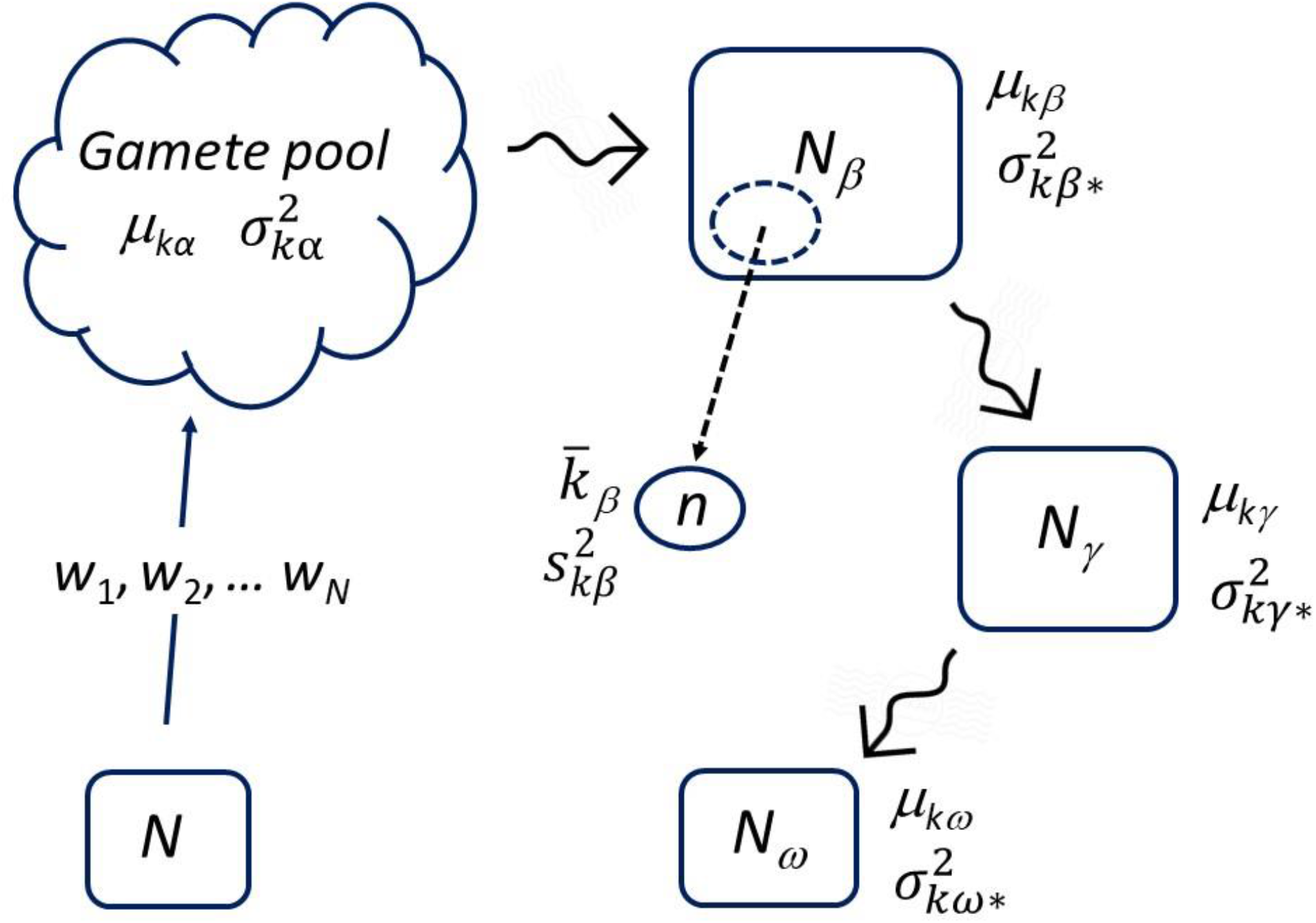
Schematic representation of a generalized Wright-Fisher model of reproduction that allows for unequal probabilities of reproductive success. *N* adults contribute gametes in the proportions *w*_1_, *w*_2_,… *w_n_* to a very large (~infinite) gamete pool (life stage α), where gametes unite at random to produce zygotes. Subsequently, random mortality (squiggly arrows) reduces the number of surviving offspring (*N_j_*) at successive life stages (*j* = *β*, γ, ω) and leads to smaller values of *μ_kj_* = mean offspring number per parent. If the number of surviving offspring at the final life stage is *N_ω_* = *N*, population size is constant population and *μ_kωj_* = 2, as in the original Wright-Fisher model. Each stage-specific variance in offspring number per parent 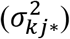 is a random variable, conditional on *μ_kj_*. Samples for empirical data analysis might be taken at any life stage (stage *β* in this example). Mean offspring number in the sample 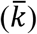 is determined by the number of offspring sampled (*n*) according to 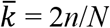; the sample variance 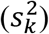 is a random variable conditional on the sample 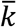 according to Equation 12.

After the gametes randomly combine to form zygotes, random survival reduces the number of surviving offspring (*N_j_*) at successive life stages. The number of life stages to consider is arbitrary; the schematic diagram in Figure 1 shows three stages following the gamete pool (*j* = α, β, γ, ω), with *j*=*ω* denoting the final (adult) stage. Mean offspring number per parent at the various life stages is determined by the stage-specific population sizes according to *μ_kj_* = 2*N_j_*/*N*. If sex ratio is equal and the number of surviving offspring at the final life stage is *N_ω_* = *N*, population size is constant and *μ_kω_* = 2, as in the original Wright-Fisher model. Each stagespecific variance in offspring number per parent 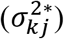 is not uniquely determined by the initial weights and *N_j_*; instead, it merely represents one of many possible random realizations of a stochastic process. In the notation, the asterisk (*) is used to denote the fact that 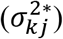 can be considered to be both a population parameter (because it applies to all individuals in the population at a given point in time) and a random variable (because it represents only one of many possible outcomes, given the fixed composition of the initial gamete pool and the survival rate through stage *j*). Waples and Faulkner (2009) also used the asterisk in a similar way in their evaluations of variation in realized reproductive parameters under the standard Wright-Fisher model.

The Crow-Morton model envisions enumerating a population at two time periods separated by random mortality, and it can be related directly to this generalized Wright-Fisher model as follows. Let time period 1 be the initial gamete pool (so time 1 = life stage α). From Equations 2–3 we have 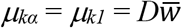 and 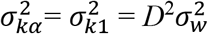, so the variance-to-mean ratio in the gamete pool is

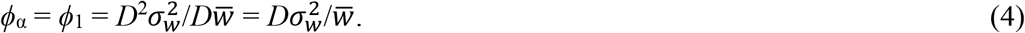

We are now interested in predicting what the realized values of 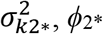, *ϕ*_2*_, and *I*_2*_ will be at a second (later) life stage, after random mortality has reduced the offspring population size to *N*_2_ and the mean offspring number to *μ_k2_* = 2*N*_2_/*N*. A simple rearrangement of Equation 1 (Waples 2002), using the notation above, leads to

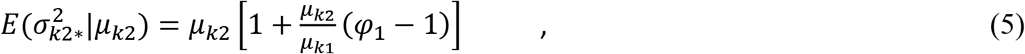

with the “|” notation indicating that this is a conditional expectation that depends on the value of *μ_k2_*. Substituting for *ϕ*_1_ from Equation 4 produces

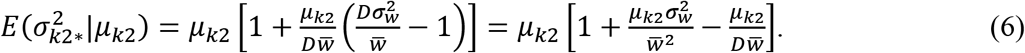

Because *D* is a very large constant, the last term can be ignored, leading to

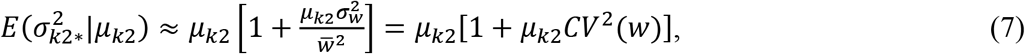

where *CV*^2^(*w*) is the squared coefficient of variation of the individual weights. Finding the conditional expectation of *ϕ*_2*_ is straightforward:

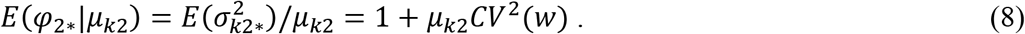

When the initial weights are equal, *CV*^2^(*w*) = 0 and we recover the standard Wright-Fisher model with *E*(*φ_drift_*) = 1. Note that this result is independent of the life stage or the stage-specific *μ_k_*, a property first demonstrated by Fisher (1939).

The analogous conditional expectation for the Opportunity for Selection is

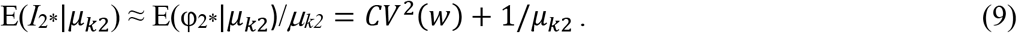

Under a pure drift model, *CV*^2^(*w*) = 0 and Equation 8 reduces to

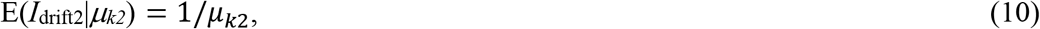

which is the inverse of the mean offspring number per parent (Downhower et al. 1987). Unlike *E*(*φ_drift_*), therefore, E(*I*_drift_) is a conditional expectation that depends on the life stage at which mean offspring number is calculated.

It is useful to define new metrics Δ_φ_ and Δ_*I*_, obtained by subtracting the random expectation from the raw value of the index. Letting subscript 1 represent raw data and subscript 2 rescaled data, we have:

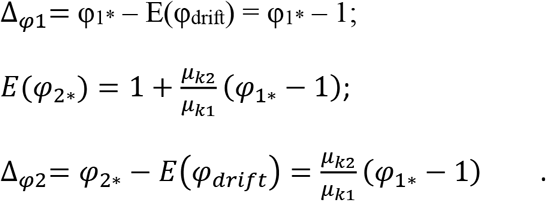

Thus, Δ_φ1_ and Δ_φ2_ differ by the factor *μ*_*k*2_/*μ*_*k*1_. Δ_φ_ quantifies the degree of overdispersion in 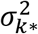, but since the adjustment only involves subtracting a constant, Δ_φ_ still depends on mean offspring number, regardless whether raw or scaled values are used.

Results are different for the Opportunity for Selection:

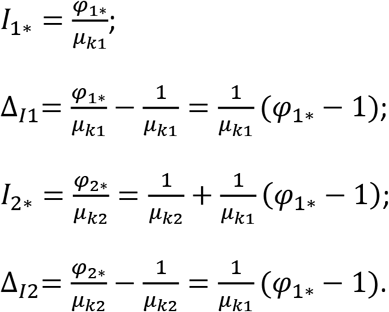

Therefore,

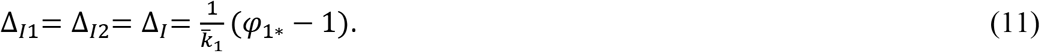

After scaling *φ*_1*_ to *μ*_*k*2_ = 2, one obtains the following:

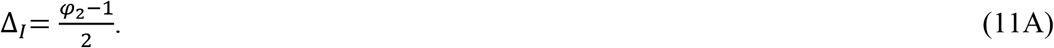

Notably, the Opportunity for Selection, adjusted to account for the contribution from random reproductive success, is the same whether it is calculated for raw or rescaled values of *I* (hence independent of mean offspring number or the life stage at which reproductive success is quantified).

Thus far we have only dealt with population parameters, including some that can be considered random variables with respect to the initial gamete pool. Samples for empirical data analysis might be taken at any life stage; in the example in Figure 1, it is stage *β*. Mean offspring number in a sample is uniquely determined by the number of offspring sampled (*n*) according to 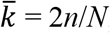, but the sample variance 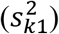 is a random realization of a stochastic process that also depends on 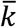. From a statistical standpoint, taking a random sample of *n* offspring from any life stage subsequent to the initial gamete pool is mathematically equivalent to finding a life stage at which random mortality has reduced the population to *N*_j_ = *n* individuals and sampling all of them. Therefore, for sampling in any life stage *j*, the expected sample variance in offspring number, conditional on mean offspring number in the sample, is given by (modified from Equation 7):

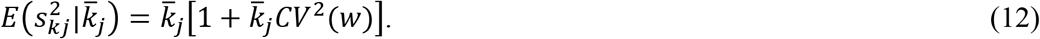

Similarly, the conditional expected values of 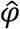 and *Î* for random samples are given by simple modifications of Equations 8 and 9:

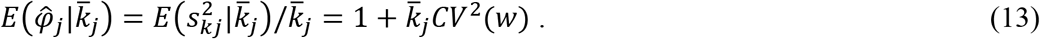

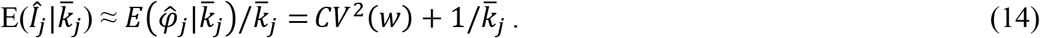

The hat “^” is used for 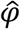 and *Î* to indicate that they are estimates based on samples (and are conditional on the sample mean offspring number) and do not reflect population parameters.

Another way to scale indices of reproductive success is to identify upper limits to possible values for 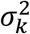, *φ*, and *I*. Ruzzante et al. (1996) provided the following expression for the maximum possible variance in offspring number among *N* parents:

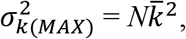

from which it is straightforward to derive the other maxima, assuming a sample of *n* offspring:

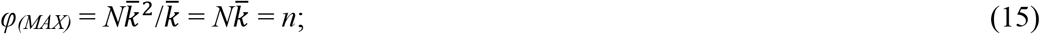

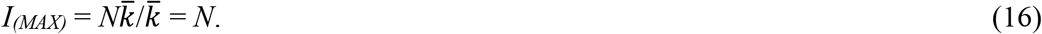

Ruzzante et al. (1996) used 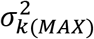 to define an index of relative resource monopolization as

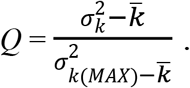

*Q* is the ratio of the empirical variance in offspring number to the maximum possible variance, after both have been adjusted by subtracting the expected value under Poisson variance in reproductive success 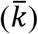. In the current notation, 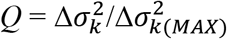. As a result of this standardization, *Q* is independent of mean reproductive success. However, as noted by Fairbairn and Wilby (2001), because *Q* is calculated using the variance in reproductive success rather than *I*, it is not readily interpretable in terms of the Opportunity for Selection. A more useful index for this purpose is

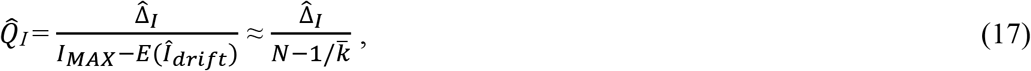

which uses *I* directly.

Crow’s Index of Variability can also be used to calculate both inbreeding and variance effective population size (Crow and Denniston 1988):

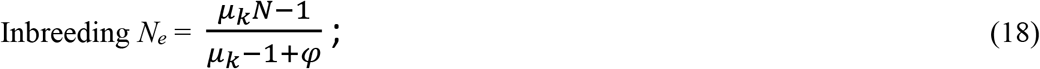

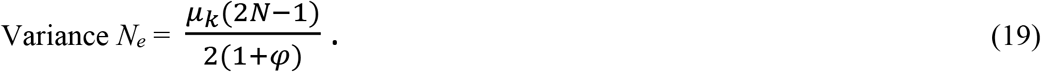

These equations give effective size for a single sex, and Eq. 19 ignores the (generally minor) adjustment for departure from Hardy-Weinberg genotypic proportions. If effective size is calculated separately for males and females (*N_em_, N_ef_*), overall *N_e_* can be obtained using Wright’s (1938) sex ratio adjustment: *N_e_* = 4*N_em_N_ef_*/(*N_em_* + *N_ef_*).

Felsenstein (2019) showed that in the weighted Wright-Fisher model, variance *N_e_* in a stable population (*u_k_* = 2) can be expressed as a simple function of the initial weights:

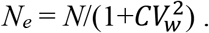

It can be shown that this result holds more generally for inbreeding effective size assessed at any given life stage *j*. Ignoring the “−1” term in the numerator, Equation 18 becomes

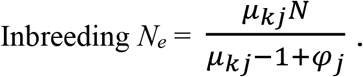

From Equation 8 we have 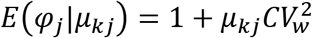, and making that substitution produces

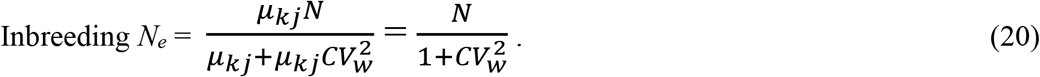

### Simulations

The analytical results above demonstrate the conditional dependence of φ and *I* on mean offspring number. To illustrate these relationships for a wide range of sample 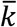 values, and to evaluate performance of the newly-proposed indices 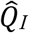 and 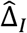, a simulation model was developed that mimics the weighted Wright-Fisher model. For each of *N_Off_* offspring a (nominally female) parent was randomly chosen from the pool of *N* potential parents. A total of *N_Off_*=100*N* offspring were produced initially, and random subsamples of different numbers of offspring (*n*) were taken to produce a series of sample 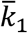 values; because only one parent was chosen for each offspring, 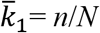. Three reproductive scenarios were considered, defined by different vectors of parental weights. (1) Each female was equally likely to be the parent of every offspring (this is the Wright-Fisher model with equal parental weights; *W*_1_ = 1, 1,… 1). (2) Moderate reproductive skew created by unequal weighting of parents (*W*_2_ = 1, 2, 3… *N*). (3) Stronger reproductive skew created by more unequal parental weights 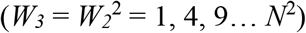. Both for the raw data and after scaling to constant population size, the statistics 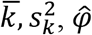, and *Î* were calculated for each sample size for each of 1,000 replicates, with 10-fold additional replication for 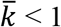. Because the ratios 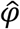 and *Î* are upwardly skewed, geometric means across replicates were used as measures of central tendency. For each sample size and resulting 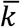, both effective sizes were estimated using geometric means of 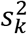 and raw and scaled 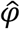 from the simulated reproductive success data.

## Results

The analytical results in **Methods** showed how expected values of the Index of Variability, the Opportunity for Selection, and effective population size can be expressed as a simple function of weighted parental contributions to an initial gamete pool. The simulation model employed a comparable scheme for assigning relative probabilities that different parents would produce offspring. Simulation results for a representative scenario with *N* = 100 parents and *N_off_* = 10^4^ total offspring are illustrated in Figures 2–4. Taking a series of random subsamples of offspring (*n* = 10 … 5000) produced results for raw 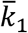 spanning three orders of magnitude (0.1 – 100).

**Figure 2.**
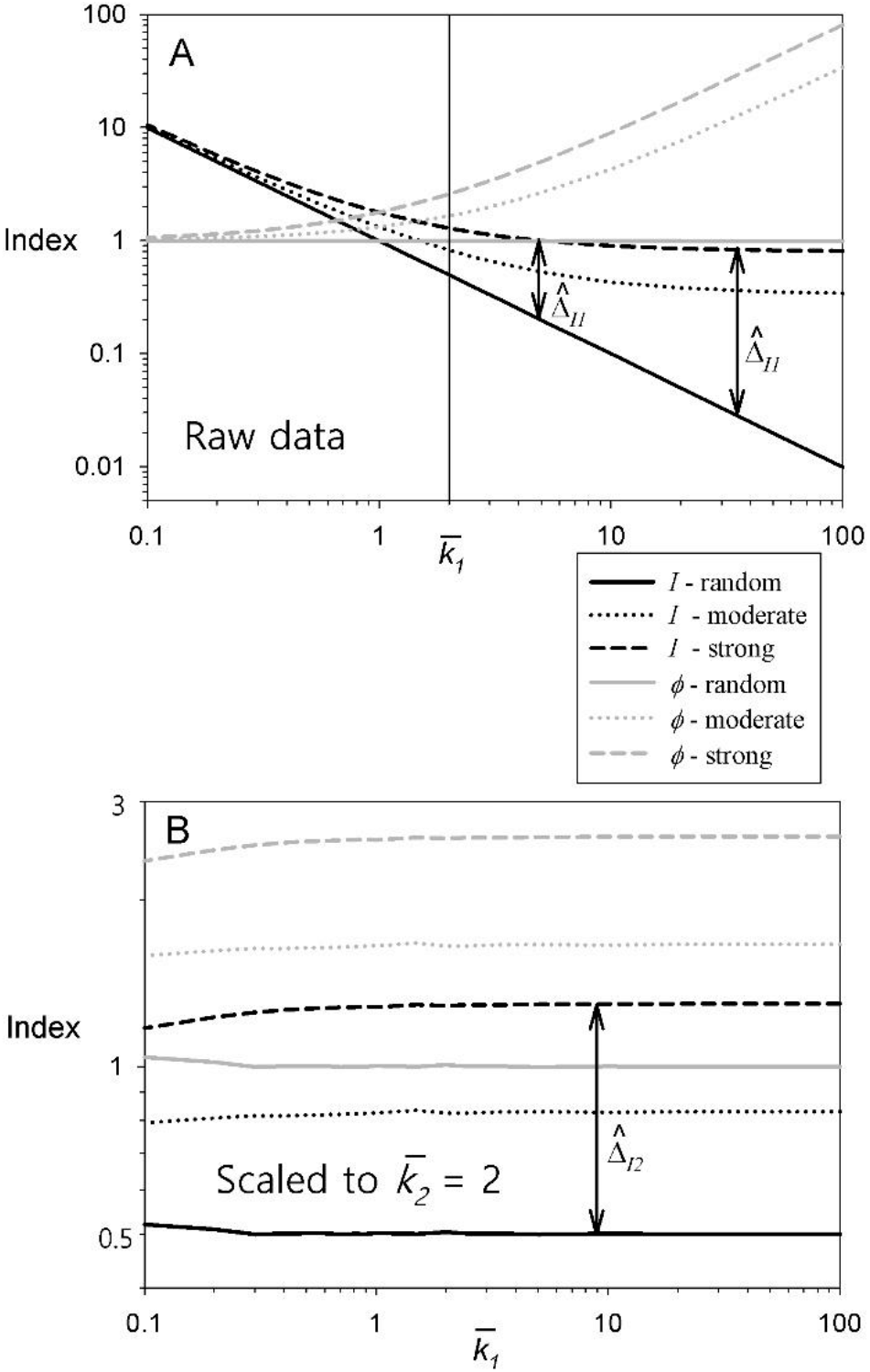
Dependence of Crow’s Index of Variability (φ) and Opportunity for Selection (*I*) on mean number of offspring per parent in the sample 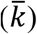. With *N* parents and *n* offspring in the sample, 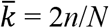. Sample 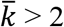 can occur in an increasing population or when sampling early life stages; 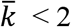 implies a declining population or incomplete sampling of offspring. Shown are geometric mean values of 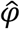, *Î*, and 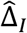 for 10,000 simulated reproductive events, using either a Wright-Fisher random reproductive success model, or moderate or strong overdispersion (unequal parental weights). A: raw data, plotted as a function of mean offspring number in the original sample 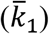. B: raw data scaled to expected values at 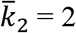, using Eq. 1. Rescaled values in panel B are essentially the same values obtained by random subsampling the original data to reach 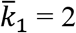 (indicated by vertical line in panel A). 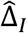 is the difference between the sample *Î* and the expected value of *Î* under random reproductive success. The three 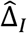 values for strong reproductive skew in the two panels (plotted at arbitrary values of 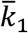) are all of the same magnitude 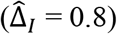.

**Figure 3.**
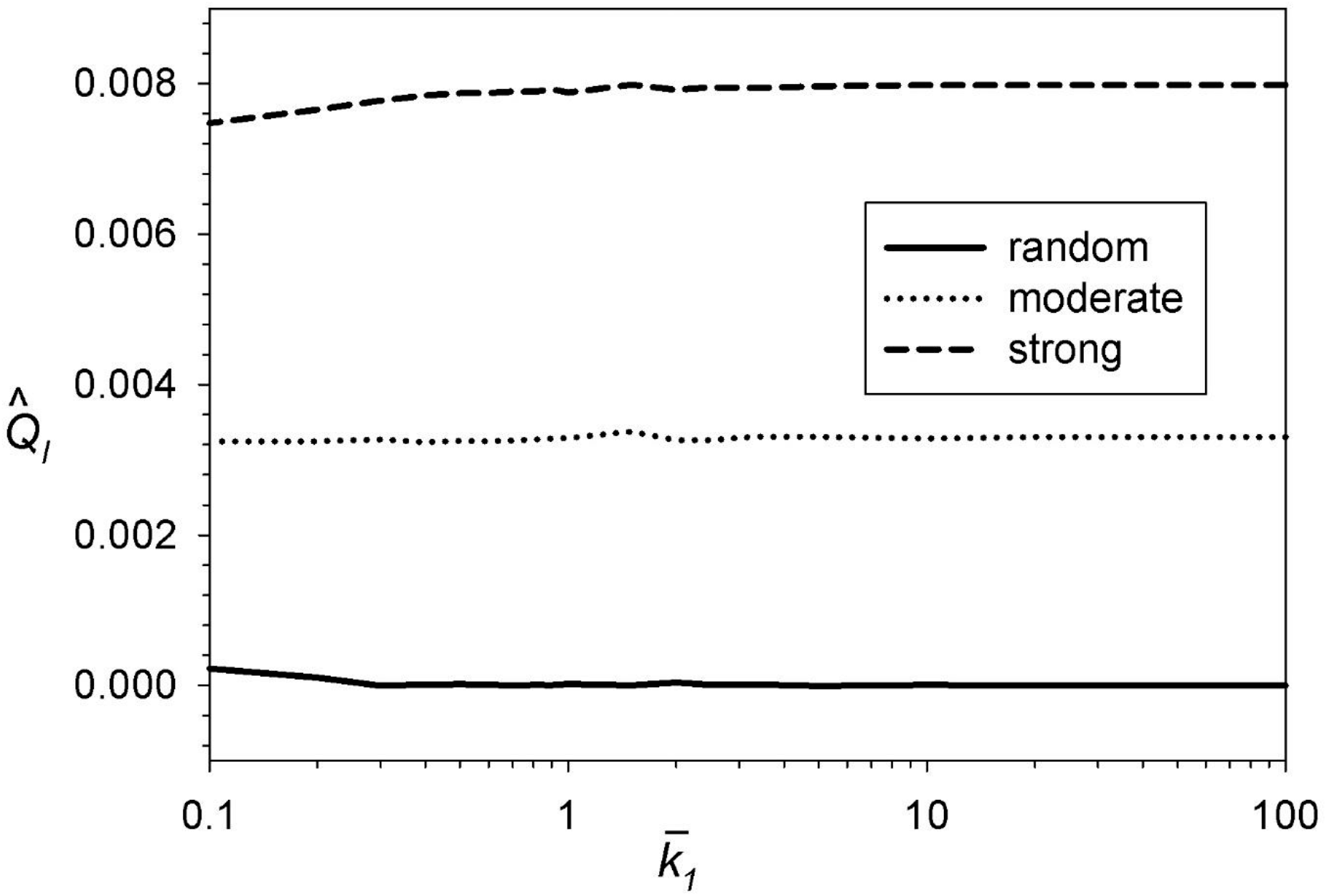
The *Q* index of relative resource monopolization, modified from Ruzzante et al. (1996) as shown in Equation 17. 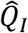 is the ratio of the empirical *Î* to the maximum possible *Î*, after both have been adjusted by subtracting the expected value under random reproductive success. Results are based on the simulated data analyzed in Figure 2.

**Figure 4.**
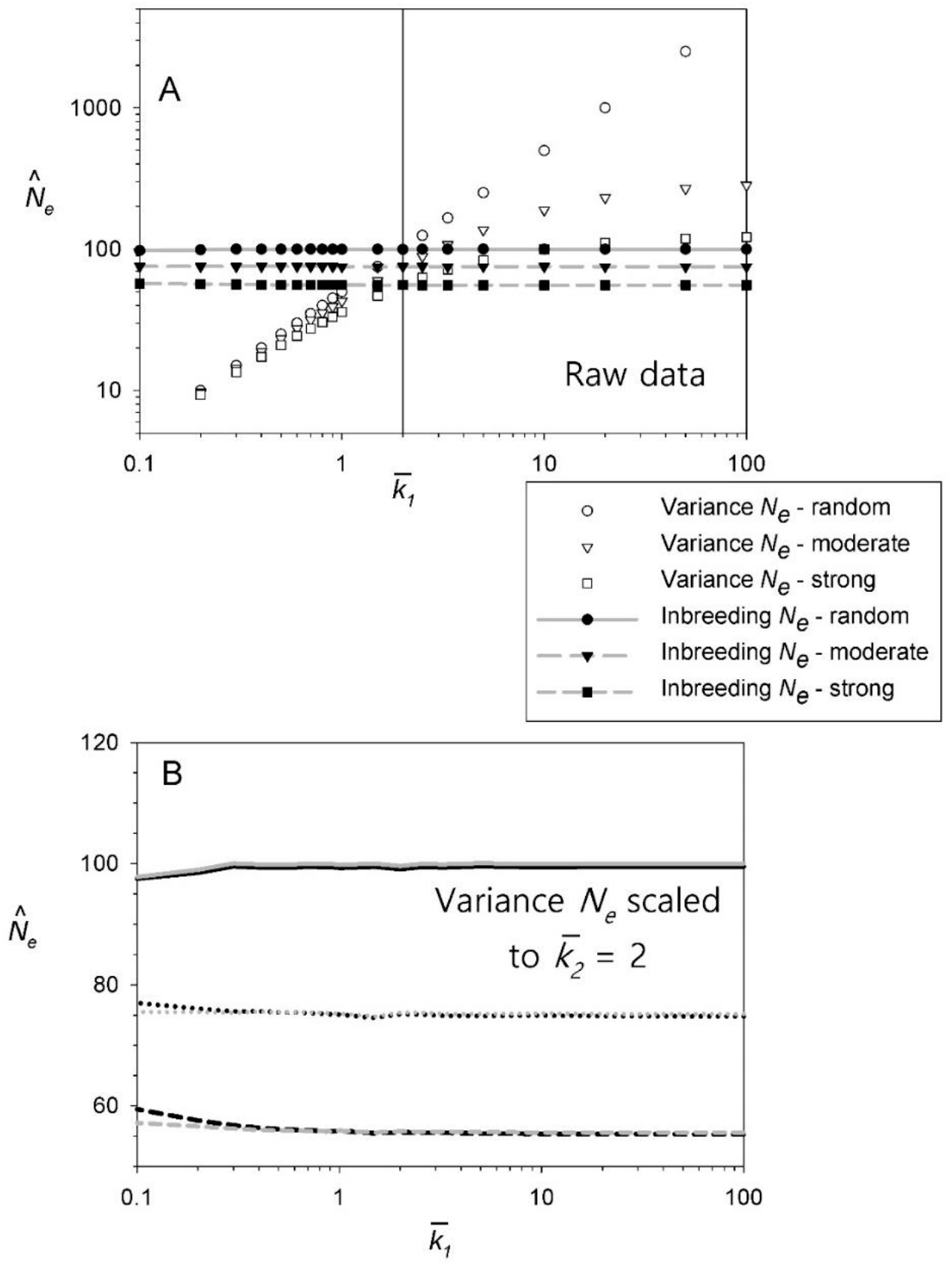
Dependence of variance 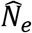 (but not inbreeding 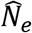) on mean offspring number in an initial sample 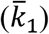, for the three reproduction scenarios considered in Figure 2 with *N* = 100 parents. A: raw data; harmonic mean variance 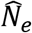 is positively correlated with 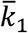 for all reproductive scenarios, whereas harmonic mean inbreeding 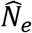 is independent of 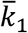. Empirical data are depicted by symbols; horizontal gray lines show true *N_e_* (from Equation 20) for the three reproductive scenarios (*N_e_* = 100 for random reproductive success, *N_e_* = 75.2 for moderate reproductive skew, and *N_e_* = 55.6 for strong reproductive skew). B: after scaling raw 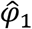 to constant population size 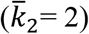 using Equation 1, variance 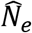 produces essentially the same result as raw (unscaled) inbreeding 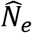. Rescaling to 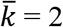 actually produces identical values for variance and inbreeding 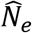; in Figure 4B, the rescaled variance 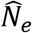 is plotted against the raw inbreeding 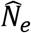 to illustrate why it is not necessary to rescale sample 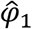 when estimating inbreeding *N_e_*.

### Crow’s indices

Sensitivity of both raw 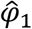 and raw *Î* to sampling intensity is apparent; the one exception is for φ under random reproductive success, in which case 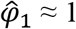 regardless what 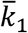 is (Figure 2A). But with overdispersed variance in reproductive success, 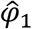 increases strongly with 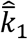 (raw 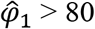 for 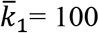 under strong reproductive skew). Underdispersed variance in reproductive success produces the mirror-image pattern: raw 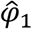 is very low (≪1) when 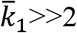 and converges on 1 as 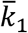 approaches 0 (Appendix A and Figure S1). It is apparent from Figure S1 that sparse sampling of offspring will generally have a poor chance of detecting underdispersed variance in reproductive success, even when underdispersion is pronounced.

For the scenarios depicted in Figure 2, stronger reproductive skew produces larger values of *Î*_1_. However, whereas 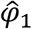 increases with 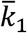 when reproductive success is overdispersed, *Î*_1_. shows the opposite pattern: the Opportunity for Selection is negatively correlated with mean offspring number (Downhower et al. 1987; Cabana and Kramer 1991).

Scaling raw 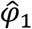 and *Î*_1_ values to a fixed 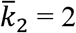 removes the dependence on mean offspring number for both indices (Figure 2B). In rescaling the empirical data for the simulation results, I used the exact expectation for random variation in offspring number (Equation 1A). Use of the Poisson approximation in Crow and Morton’s formula (Equation 1) produces a slight bias when 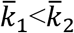 (Figure S2). The scaled values in Figure 2B, obtained from the raw values in Figure 1A using Equation 1, are the same values identified in Figure 2A by the intersection of the vertical line for 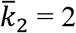 and the curves for the empirical data. That is, subsampling from the original data to reach 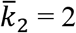 produces essentially the same result as scaling the raw data using the Crow and Morton method. Note, however, that for both 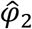 and *Î*_2_, the magnitude of the rescaled values depends on choice of 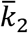.

Independence of 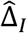 from both raw and scaled 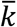 is also illustrated in Figure 2. Although it is not visually apparent with the log scaling of the *Y* axis, for strong reproductive skew the two 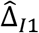 values for different 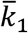 Figure 2A are the same 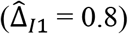, and both are equal to rescaled 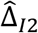 from Figure 2B (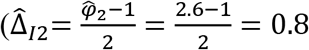 using Equation 11A).

### Index of resource monopolization

In the simulations, the estimator 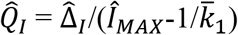 (modified from Ruzzante et al. 1996 according to Eq. 17) was independent of sample 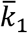 under both random and overdispersed variance in reproductive success (Figure 3). Equations 15 and 16 show that maximum possible 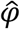 is just the number of offspring sampled (*n*), whereas the maximum possible *Î* is the number of parents. These results show that 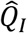 could be useful in quantifying the effects of resource monopolization in the context of the Opportunity for Selection (Ruzzante et al. 1996; Fairbairn and Wilby 2001).

### Effective population size

With random reproductive success, E(φ) ≈ 1 and inbreeding *N_e_* ≈ *μ_k_N*/*μ_k_* = *N*, regardless what *μ_k_* is because inbreeding *N_e_* depends on the number of parents (Crow 1954; Equation 18). In contrast, for variance *N_e_*, the *μ_k_* term in the numerator is not balanced by a separate *μ_k_* term in the denominator (Equation 19), and as a consequence variance *N_e_* depends heavily on mean offspring number in a sample (Figure 4A). Harmonic mean inbreeding 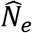 is invariant with 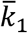 for all reproductive scenarios, and the sample estimates agree almost exactly with the true *N_e_* predicted by Equation 20 based on weighted contributions to the initial gamete pool. In contrast, variance 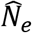 is directly proportional to 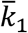 for random reproductive success and has a more complex but positive correlation with 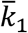 when variance in reproductive success is overdispersed. The two effective sizes are the same only when 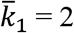 (vertical line in Figure 4A), indicative of a stable population (Crow 1954). Rescaling 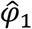 to its expectation for a population of constant size removes the dependence of variance 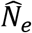 on mean offspring number (Figure 4B).

### Effects of age structure

All of the above analyses have implicitly assumed that generations are discrete. Because most species are age structured (Hughes 2017), this is an important limitation. In age-structured species having birth-pulse reproduction (Caswell 2001), there are two major ways to calculate the mean and variance in reproductive success: 1) seasonal reproductive success among all mature individuals during a single year or season; 2) lifetime reproductive success among all individuals from the same cohort. Below I consider some of the consequences of age structure for Crow’s indices of reproductive success and related analyses.

#### Index of Variability

Analysis of seasonal reproductive success conforms to the discrete-generation model, with one exception: reproductive success for all adults in a single season can be partitioned by parental age. Seasonal reproduction within an age-structured population can be described by vectors of age-specific mean reproductive success and age-specific variance in reproductive success: 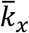 and 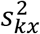, where *x* indicates age. 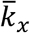 is an estimate of the familiar *b_x_* (or *m_x_*) from a standard life table, published data for which are available for many species. 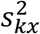 is the analogous age-specific sample variance, but empirical estimates of this key parameter are seldom published. The ratio of the age-specific variance and mean is 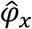 (Waples et al. 2011; Waples 2016), which also is rarely reported in the literature.

As is the case with discrete generations, 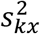 and 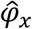 are sensitive to 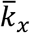. These indices can be rescaled to a different mean offspring number using Equation 1, but there is a complication: it is not generally the case that scaling to 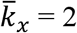 will produce results expected for a stable population. Instead, 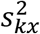 and 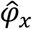 can be rescaled to the vector of *b_x_* values that produce a stable population. This requires that measures of relative, age-specific reproductive success be converted to measures of absolute reproductive success by ensuring that Σ*l_x_b_x_* = 2, where *l_x_* is cumulative survival through age *x*. This in turn requires knowing or being able to estimate age-specific survival rates. Table 2 in Waples et al. (2018) shows how age-specific 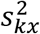 and 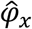 values can be rescaled to constant size in a species with overlapping generations.

#### Effective size

Because of its insensitivity to mean offspring number, inbreeding effective size is much more generally useful than is variance effective size when analyzing age-structured data. For these species, Equation 18 can be used to estimate the effective number of breeders per year or season (*N_b_*) (Waples 1990; 2016), which generally differs from *N_e_* per generation. For this purpose, the age-specific vectors of 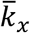 and 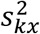 values are converted into scalars 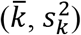 that reflect seasonal reproductive success for adults of all ages. Estimating *N_e_* per generation requires integration of reproductive success data across multiple years or seasons to obtain the lifetime estimators 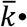 and 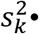; see Waples et al. (2011, 2014) for details.

#### Opportunity for Selection

Although the concept of age-specific Opportunity for Selection isn’t particularly meaningful, Crow’s *I* can be (and often is) calculated in two fundamentally different ways: (1) among all individuals reproducing in a single year or season, and (2) using lifetime reproductive output among all individuals in a single cohort. For seasonal data, 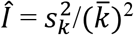 and 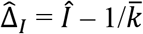. As is the case with discrete-generation data, 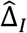 for seasonal reproduction is independent of mean offspring number in the sample, whether raw or scaled data are used.

When applied to lifetime reproductive success data for members of a cohort, 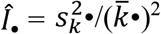 is appropriate when estimating the Opportunity for Selection. Calculation of E(*Î*_•drift_) is complicated by the necessity of considering random stochasticity in both survival and reproduction over individual lifetimes. In general, even under a null model that assumes all individuals in a cohort experience the same age-specific probabilities of survival and reproduction, lifetime 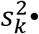 will exceed 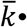 because some individuals, by chance, survive longer and have more opportunities to reproduce. It does not appear that an analytical approximation is currently available for lifetime E(*Î*_•drift_). However, under the null model, a stochastic, absorbing Markov chain model with rewards developed by Caswell and colleagues (Caswell 2011; van Daalen and Caswell 2017) could be used to calculate lifetime E(*Î*_•drift_) and hence lifetime 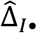, given information on age-specific vital rates from a standard life table. A limitation of the Caswell model is that it implicitly assumes that φ_x_ ≈ 1 for each age and sex. The *AgeNe* model (Waples et al. 2011) is more general in this respect, as it allows for age-specific and sex-specific values of φ_x_, but has the limitation that it is deterministic. Age-structured simulations have shown, however, that *AgeNe* accurately predicts lifetime 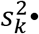 under conditions of random demographic stochasticity, as envisioned by Caswell and colleagues (Waples et al. 2014). Therefore, lifetime 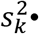, calculated in *AgeNe* after setting φ_x_ = 1 and assuming lifetime 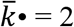, could be used to estimate 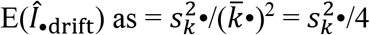. From this one could estimate lifetime Δ_I_ as 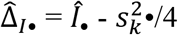.

## Discussion

The weighted Wright-Fisher model used here is a convenient way to link the analytical and simulation models, which produced congruent results. This model provides a way to express the Index of Variability, the Opportunity for Selection, and effective population size as simple functions of proportional contributions by individual parents to an initial gamete pool. Major points to emerge include the following:

1. Rescaling *Î* to a fixed 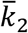 (as Crow and Morton proposed for the Index of Variability many decades ago) removes the dependence of the Opportunity for Selection on mean offspring number, but the result still depends on choice of 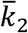. Although an argument can be made that 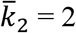 is a logical reference point for standardizing offspring number, that is not the only plausible target for variance rescaling. Furthermore, this transformation raises the question whether rescaled *Î* can still be interpreted as the variance in relative fitness.
2. The new index 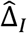 introduced here solves this problem by simultaneously (1) removing the dependence of *Î* on mean fitness in the sample, without the need for rescaling variance in reproductive success; and (2) quantifying the degree to which empirical *Î* exceeds the value expected under a null model of random variation in reproductive success. This means that Δ_*I*_ can be used to compare empirical estimates of the Opportunity for Selection from:

- different studies;
- different samples within the same study;
- samples taken from different life stages;
- samples of males and females when the sex ratio is uneven. The 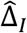 index also directly addresses the need for an appropriate null model for the Opportunity for Selection, which has been pointed out by numerous authors (e.g., McClain 1986; Downhower et al. 1987; Ruzzante et al. 1996). Although the analytical and numerical methods used here modeled diploids with separate sexes, the approach could easily be extended to monoecious or haploid populations.
3. The modified index *Q_I_* is cast in terms of the Opportunity for Selection; it is independent of mean offspring number and might have some practical utility. However, conditions that lead to *I_MAX_*, where one or a few parents produce all the offspring and the rest of the parents none, are quite extreme for any stable population. This is the type of scenario proposed by Hedgecock (1994) in his Sweepstakes Reproductive Success hypothesis, which might apply to some marine fishes with high fecundity and Type III survivorship (Hauser and Carvalho 2008; Hedgecock and Pudovkin 2011). For most species, however, scaling *Q* using *I_MAX_* will generally produce very low 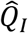 values even when overdispersion is relatively strong (Figure 3).
4. Simulation results confirm that Equation 20 provides a simple way to predict inbreeding effective size based on individual weights that reflect relative probabilities that different individuals will be the parent of a random offspring. With modern genomics technology and access to methods that can non-invasively sample DNA from individuals in the wild, parentage analyses are now routinely conducted for natural populations of many plants and animals. Such data are typically collected annually and later integrated across years. Each set of annual reproductive success data can be used to estimate the annual effective number of breeders, *N_b_*, and the ratio *N_b_*/*N*. These demographic estimates can also be compared with single-sample genetic estimates of inbreeding *N_b_*, which are now widely applied to natural populations (Palstra and Fraser 2012; Whitely et al. 2015; Wang 2016). Because of its sensitivity to sample 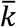, variance effective size is not well-suited for these applications.

A case can be made that Crow’s *I* can be meaningfully applied to either seasonal or lifetime reproductive success data, and examples of both are common in the literature. Quantifying variance in lifetime reproductive success is clearly important, as this parameter strongly influences *Ne* per generation and long-term evolutionary behavior of the population. Calculation of 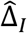 based on lifetime reproductive success is complicated by having to account for stochastisity in both survival and reproduction propagated across the lifespan (Brommer et al. 2002), but the stochastic “Markov chain with rewards” model of Caswell and colleagues should be useful here. On the other hand, reproduction only occurs among individuals that co-occur in space and time. Furthermore, sexual selection in terms of competition for mates only occurs among individuals reproducing in the same season. Seasonal/annual data on variance in reproductive success and the estimators 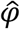 and *Î* therefore can provide valuable insights into mating systems and behavioral dynamics associated with reproduction.

Although Δ_*I*_ provides a simple way to standardize empirical estimates of Crow’s *I*, interpretation of the Opportunity for Selection remains complicated. Even if 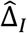 is positive, it does not prove that natural selection is operating–it only shows that the distribution of reproductive success is more skewed than expected under random mating dynamics. As an alternative to random survival, Crow and Morton (1955) considered an extreme case of family correlated mortality, in which entire families either survive or do not as a unit. This type of result could occur in two general ways. First, survival might reflect individual survival phenotypes of the offspring or nurturing phenotypes of the parents. In this scenario, high mortality within a clutch of offspring could be due to failure of the offspring to grow quickly and avoid predators, or failure of the parent to provide food or ward off predators. To the extent that these phenotypes have a genetic basis, this would reflect the operation of natural selection. This can be called the “bad genes” explanation for family correlated mortality.

Under an alternative scenario, most or all offspring from a family might fail to survive merely because, for a short period of time, they share a common environment. This could occur, for example, if the tree supporting a bird nest blows over in a storm, or if all the eggs a female salmon lays in the gravel get washed away during a flood. In these scenarios, survival or mortality might have nothing to do with offspring phenotypes or genotypes; instead, it could simply be a case of being in the wrong place at the wrong time, together with your siblings. This can be called the “bad luck” explanation. Parsing family-correlated survival into “bad genes” vs “bad luck” hypotheses can be complicated and is likely to be species and population specific (e.g., see Snyder and Ellner 2018). One needs to consider questions such as, “When choosing nest sites, would some birds (those with better genes) have avoided trees likely to suffer storm damage?” and “Would a bigger, stronger female salmon have dug a nest deep enough to withstand high flows, or located the nest in an area less prone to scouring?” This and related issues are why most authors refer to Crow’s *I* as the Opportunity for Selection; his original term Index of Total Selection seems to imply a more direct link to actual natural selection.

One potential caveat applies to use of Δ_*I*_. This index is most useful when, as will often be the case, mean offspring number in a sample depends on logistical constraints or aspects of experimental design such as sampling intensity or life stage sampled. In these situations, sample 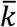 largely represents noise in the analysis and its effects are best removed. However, mean fitness in the population as a whole (*u_k_*) is a key parameter that can reflect differential response to selection under different environmental conditions (e.g, Cao et al. 2019). In cases where it is possible to essentially inventory the entire population (such that 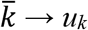), applying the Δ_*i*_ adjustment could be counterproductive by removing part of the signal.

### Sampling considerations

The above analyses have assumed random subsampling of all offspring. Family-correlated sampling, in which siblings are collected together more often than would occur by chance, can affect the estimate of mean fitness and can have complex, cascading effects on 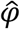 and *Î*. These sampling effects can be difficult to avoid for highly fecund species when offspring are sampled at an early life stage.

For adults, a key consideration is whether sampling probability is independent of (a) an individual’s reproductive success, and (b) whether its offspring also have been sampled. If both criteria are met, then randomly sampling just a portion of the adults should not lead to bias, although it would increase uncertainty in extrapolating from the sample to the population as a whole. Also, performance of some genetic parentage programs can degrade if the fraction of unsampled parents becomes too large (Jones et al. 2010), so incomplete sampling of potential parents could indirectly affect performance of indices that estimate variance of reproductive success.

Probably the most pervasive source of potential bias in sampling adults involves those that produce no offspring. Under random reproductive success, it is expected that some fraction of mature adults will produce no offspring in a given year/season, or even across their entire lifetime; with overdispersed variance in reproductive success, the fraction of null parents will be higher. In practical applications, it is essential to know whether these null parents and those that actually produce offspring are equally likely to be sampled. This equiprobable-sampling criterion would be violated, for example, if potential parents are only sampled on the breeding grounds, but each year some fraction of adults skip breeding. Failing to account for these null parents would overestimate mean fitness and underestimate variance in reproductive success and related quantities (Hadfield 2007; Klug et al. 2010; Waples and Antao 2014; van Daalen and Caswell 2019). Interestingly, failing to sample non-reproducers has no effect on inbreeding 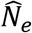; this type of sampling would affect all the parameters in Equation 12, but they change in such a way that 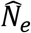 and 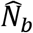 remain constant (Waples and Waples 2011). However, this scenario would affect the estimated *N_e_*/*N* or *N_b_*/*N* ratios, if some fraction of mature adults are not accounted for in the denominator.

**Table 1.**
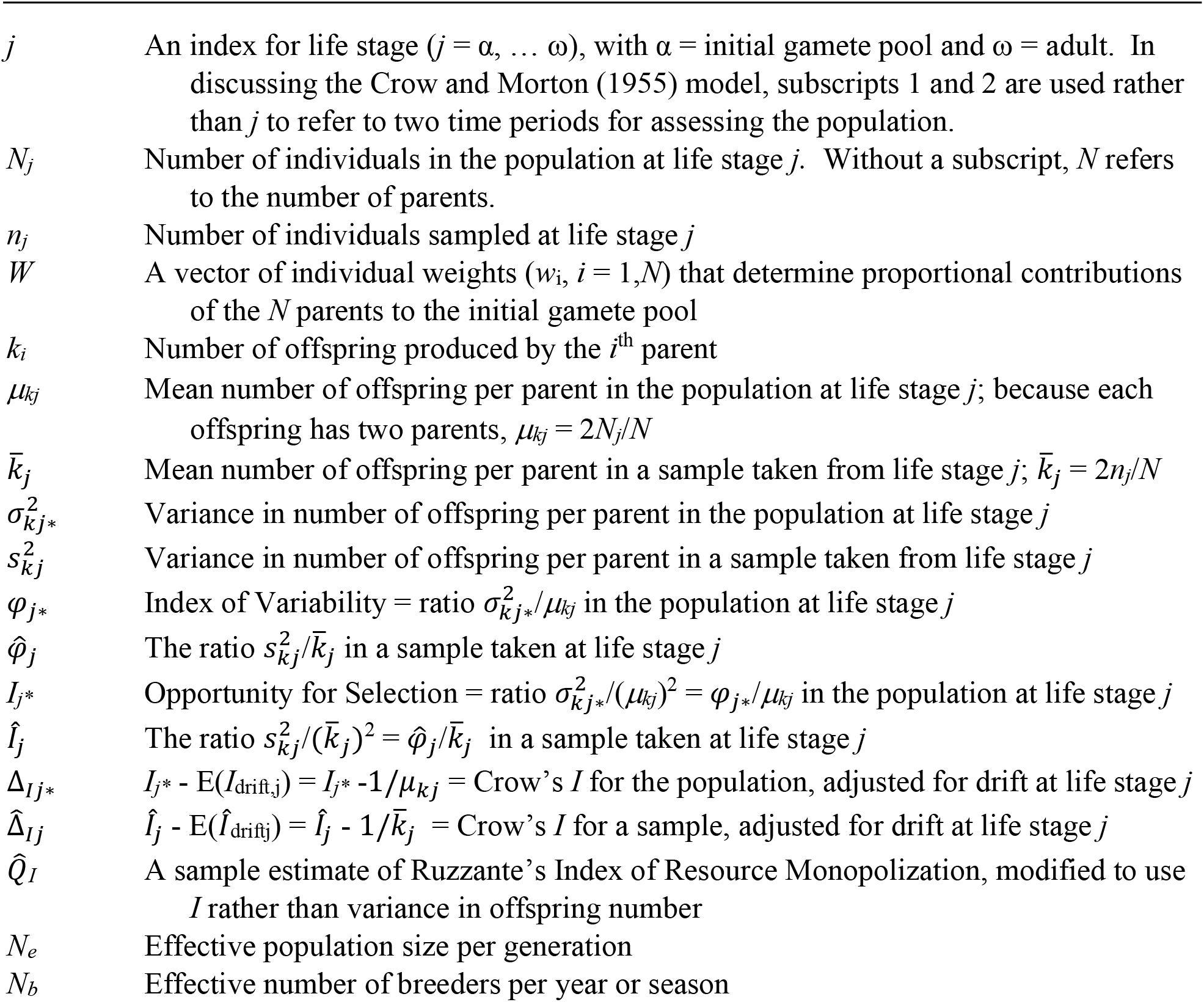
Notation used in this document. In some instances, not all of the subscripts are used. Asterisks (*) indicate terms that are both population parameters and random variables whose expectations depend on a stochastic process of random sampling from the initial gamete pool.

## Acknowledgments

I am grateful to Joe Felsenstein, Marty Kardos, and Tom Reed for useful discussions and comments. Daniel Ruzzante, Jarle Tufto, and an anonymous reviewer provided useful comments on an earlier draft. The author declares no conflict of interest.

## Appendix

## A. Underdispersed variance in reproductive success

This paper has focused on overdispered variance in reproductive success (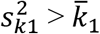, using the Poisson approximation), which reduces effective size and provides opportunities for natural selection to operate. Underdispersion 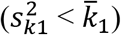 is less common but nevertheless occurs in many taxa, at least with respect to seasonal reproduction. For example, the number of eggs or early-juvenile offspring can be constrained to a relatively small range of values, and this limits how large 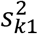 can be. At the extreme, females of many large animal species, and some birds, can only produce 0 or 1 offspring per season, in which case it is impossible for the variance in offspring number to be as large as the mean. This can be shown as follows. Of the *N* females, assume that *XN* produce exactly 1 offspring that appears in a sample, and the rest produce none. Mean offspring number in the sample is 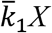 and 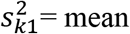 of the squares minus the square of the 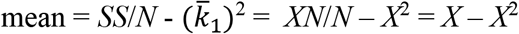, which leads to 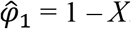.

The original Wright-Fisher model of reproduction incorporated an extreme version of this underdispersion scenario: each of *N* potential parents were imagined to contribute equally to an infinitely large pool of gametes, which then united at random to form the next generation of *N* individuals. Under this scenario, using the current notation, 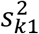 and 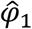 were both 0 while 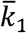 was infinitely large, and substituting these values into Eq. 1A produces

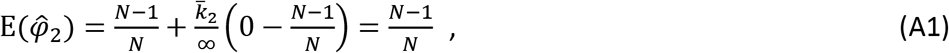

which is simply 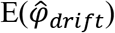 (i.e., RS is neither overdispersed nor underdispersed). Note that because the initial gamete pool (and hence 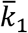) is assumed to be infinitely large, this result holds regardless what 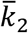 is.

All real populations of are of course finite in size. Also, we can generalize the idea of an initial gamete pool to production of diploid offspring ranging from zygotes to adults. If we still assume equal contributions to the initial pool of offspring 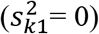, but in finite amounts by each parent, we have

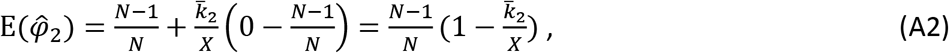

where 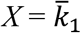 could be relatively large but is finite. Equation A2 shows that 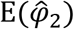 is always less than the random expectation, so variance in RS is underdispersed, and the degree of underdispersion depends on the ratio 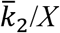.

This model is still not very realistic, as it assumes 0 variance among parents in contributions to the initial gamete pool. Overdispersion can be generated by having unequal parental contributions to the initial gamete pool. Waples et al. (2018) used a variation of this approach to provide a minimum estimate of 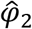 in southern bluefin tuna (*Thunnus maccoyii*), based on empirical data for variation in female size at specific ages (and hence assumed variance in age-specific egg production). For the current project, in modeling underdispersion I relaxed that restrictive assumption that 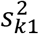 0 by having two separate processes for producing offspring: 1) a directed process, whereby each parent contributes equally to the fraction *Y* of all offspring; and 2) a random process, whereby parents for the remaining fraction (1-*Y*) of all offspring are chosen randomly from the *N* candidates. As in the main text, in the simulations 100 parents produced a total of 10,000 initial offspring (so 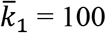), and random subsampling produced 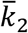 values as small as 0.1. I considered two values of *Y*: 0.5 (moderate underdispersion) and 0.9 (strong underdispersion).

Results (Figure S1) show that initial Index of Variability for the two scenarios is simply the fraction of offspring that were randomly assigned to parents (i.e., 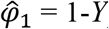). As the sample size of offspring declines, 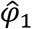 for both scenarios rapidly approach 1, in agreement with predictions from Equation 1A. It is apparent that sparse sampling of offspring will generally have a poor chance of detecting underdispersed variance in reproductive success, even when underdispersion is pronounced.

**Figure S1.**
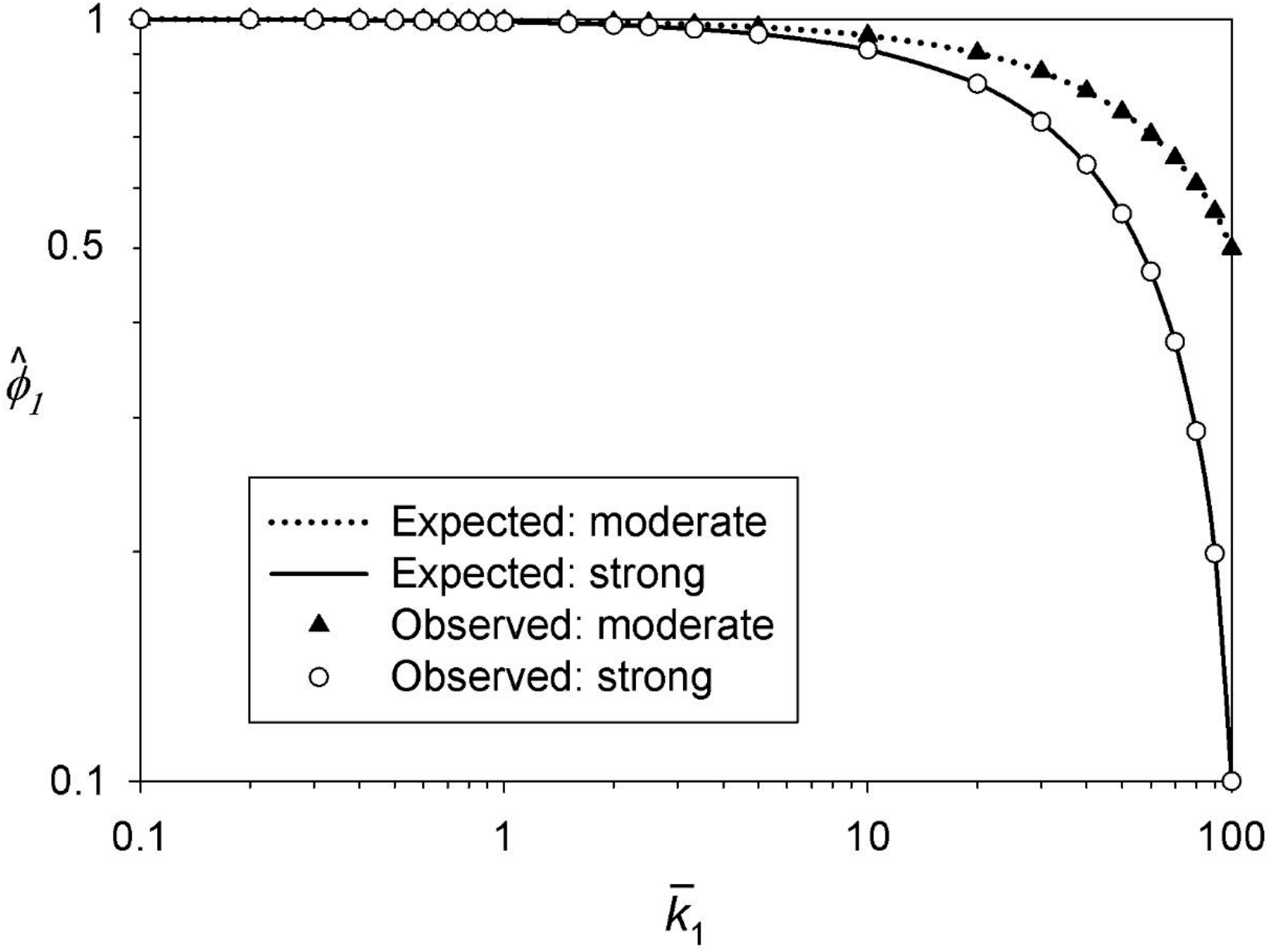
Relationship between the estimated Index of Variability 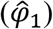 and sample 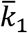 when variance in reproductive success is underdispersed. Observed values (symbols) are from simulations that initially generated 10,000 offspring from 100 parents and then randomly subsampled offspring to produce smaller sample 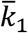 values. Expected values (lines) were obtained from Equation 1A using the empirical value of raw 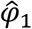 for maximum 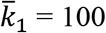. Moderate and strong underdispersion scenarios are described in the Appendix text.

## B. Poisson approximation to the binomial variance

**Figure S2.**
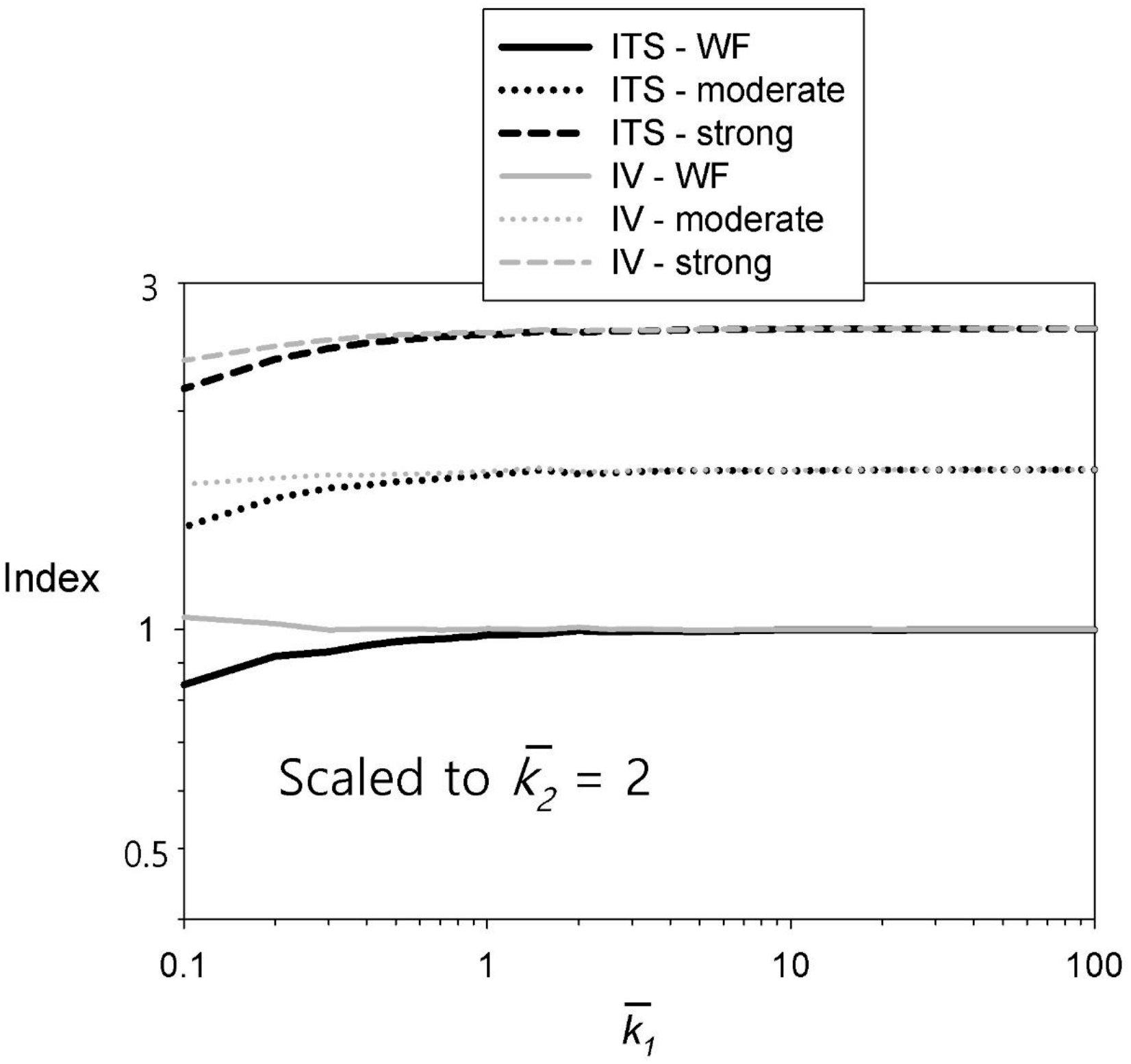
Comparison of results for rescaling empirical data for 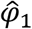 from Figure 2A to 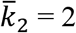, using Equation 1 (black) and Equation 1A (gray). The two equations differ in that 1 uses the Poisson approximation that 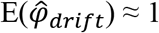, whereas 1A uses the exact value 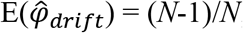, with *N* being the number of parents. The difference between (*N*-1)/*N* and 1 is small (0.99 vs 1 for the *N* = 100 parents used in simulating the data), but that difference becomes magnified when 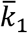 is small, in which case the term 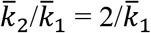 becomes large.

